# OsNLP4-OsD3 module integrates nitrogen-iron nutrient signals to promote rice tillering by repressing strigolactone signaling

**DOI:** 10.1101/2023.02.28.530551

**Authors:** Ying Song, Guang-Yu Wan, Jing-Xian Wang, Lin-Hui Yu, Jie Wu, Cheng-Bin Xiang

## Abstract

Rice tillers are a major yield component regulated by phytohormones and nutrients. How nutrients interact with phytohormones to control tillering remains largely elusive. Here, we report a novel mechanism by which the transcription factor NIN-like protein 4 (OsNLP4) integrates nitrogen (N)-iron (Fe) nutrient signals to promote tillering by repressing *OsD3* in strigolactone (SL) signaling. We show that the N-Fe balance modulates OsNLP4 nuclear accumulation, which is increased by Fe through H_2_O_2_ reduction. Furthermore, OsNLP4 upregulates multiple H_2_O_2_ scavenging genes, providing a positive regulatory loop for OsNLP4 nuclear accumulation. Our findings uncover a fundamental mechanism by which the OsNLP4-OsD3 module integrates N-Fe nutrient signals to downregulate SL signaling and thereby promote rice tillering and yield, thus facilitating sustainable agriculture worldwide.

## Introduction

Ensuring global food security is quite urgent, as the current rates of increase in crop yields are not enough to meet the 60% rise in demand by 2050 (Long et al., 2015). To feed the rapidly increasing global population, we must boost crop yield in an environment-friendly and sustainable manner (Horton et al., 2021). Rice tillers are specialized grain-bearing organs with great agricultural significance that directly contribute to grain yield (Wang et al., 2022). Tillering is controlled by multiple factors, including hormones and nutrients, which have been extensively studied in recent decades with significant progress (Wang et al., 2018). However, how nutrient signals integrate with hormone pathways to coregulate tillering is much less known. Little is known about how interactions between nutrients affect tillering.

Auxin and cytokinin (CK) are two phytohormones that regulate tillering in an antagonistic manner, with auxin inhibiting and CK promoting tillering (Wang and Li, 2011). Auxin signaling regulates tillers in an auxin transport canalization-based pathway or through second messengers such as CK (Muller and Leyser, 2011). Endogenous CKs can be transported apically through xylem sap into axillary buds and promote their outgrowth to accelerate tillering (Muller and Leyser, 2011). Gibberellin (GA) and brassinosteroid (BR) have also been reported to affect tillering. GAs inhibit tillering by triggering DELLA protein SLENDER RICE 1 (SLR1) degradation to cause MONOCULM 1 (OsMOC1) catabolism or by positively regulating the expression of rice *HOMEOBOX 1* (*OsOSH1*) and *TEOSINTE BRANCHED1/FINE CULM1* (Os*TB1/FC1*) (Liao et al., 2019). BR signaling strongly promotes tillering through the BRASSINAZOLE-RESISTANT 1 (OsBZR1)-REDUCED LEAF ANGLE 1 (RLA1)-DWARF AND LOW TILLERING (DLT) module, inhibiting *OsFC1* expression (Fang et al., 2020). In addition, moderately enhanced abscisic acid (ABA) maintains tillering by antagonizing the OsMOC1 degradation pathway promoted by GA-APC/C^TE^ (Lin et al., 2020).

Strigolactones (SLs) are a new class of carotenoid-derived phytohormones that suppress shoot branching (tillering) by inhibiting axillary bud growth (Wang et al., 2020). A series of rice *dwarf* (*d*) mutants producing a large number of tillers have been found to be involved in SL biosynthesis and signaling, confirming the unique role of SL in rice tillering (Jiang et al., 2013; Zhou et al., 2013). OsD14, OsD3 and OsD53 are core components of SL signaling. In the presence of SL, the SL receptor OsD14 binds to the F-box component OsD3 of the SKP-Cullin-F-box (SCF) E3 ubiquitin ligase complex to form OsD14-SCF^OsD3^ to target and degrade the SL repressor OsD53, thus realizing SL signal transduction (Jiang *et al*., 2013; Zhou *et al*., 2013). In the absence of SL, OsD53 promotes tillering by interacting with TOPLESS (OsTPL/TRP) and OsBZR1 to inhibit the tillering-suppressive genes *OsFC1* and *CYTOKININ OXIDASE/DEHYDROGENASE 9* (*OsCKX9*) (Duan et al., 2019; Fang *et al*., 2020).

In addition to phytohormones, the impact of nutrients on rice tillering is also well known, especially the essential macronutrient nitrogen (N). Rice tillering can be significantly affected by N supply (Wang *et al*., 2018). NITROGEN-MEDIATED TILLER GROWTH RESPONSE 5 (OsNGR5) promotes tillering by interacting with POLYCOMB REPRESSIVE COMPLEX 2 (OsPRC2) to methylate and inhibit tillering suppressors in an N-dependent manner (Wu et al., 2020). TEOSINTE BRANCHED/CYCLOIDEA/PCF 19 (OsTCP19) conveys the regulation of tillering by N through its transcriptional response to N and targets the tillering-stimulating gene *OsDLT* (Liu et al., 2021).

The micronutrient Fe is an essential component of N assimilation enzymes, electron transport of photosynthesis in the form of Fe-S clusters (Briat et al., 2015), and key enzymes in SL synthesis, such as OsD27 (Lin et al., 2009), which may indirectly affect tillering. Nutrient balance may be one of the most important factors affecting tillering due to synergistic or antagonistic effects between plant nutrients. A relative balance between N and phosphorus or potassium improved rice growth and yield, accompanied by a significant increase in tiller number (Fageria and Oliveira, 2014; Hu et al., 2019; Zhang et al., 2010). However, the effect of the N and Fe balance on tillering has not been reported to date.

We previously reported that rice NIN-like proteins (OsNLPs) are key nitrate-responsive transcription factors in N metabolism that promote N uptake and utilization (Alfatih et al., 2020; Wu et al., 2021; Zhang et al., 2022). We have continued to unravel new roles of OsNLPs and further demonstrated that OsNLP4 is a pivotal regulator of the N-Fe balance in maximizing rice yield and NUE by promoting tillering (companion paper). In this study, we specifically address the molecular mechanism of how the N-Fe balance controls tillering in rice through OsNLP4. From our RNA-seq analyses, we found that the transcript levels of *OsD3, OsCKX9* and *OsFC1* involved in SL signaling and tillering suppression were significantly altered. Subsequently, we demonstrated that OsNLP4 directly represses *OsD3* expression and thereby downregulates SL signaling, leading to increased tiller number. The N-Fe balance modulates OsNLP4 levels in the nucleus by affecting H_2_O_2_ levels, which are negatively associated with OsNLP4 nuclear accumulation. Fe supply decreases the H_2_O_2_ level and increases the nuclear accumulation of OsNLP4. In addition, OsNLP4 upregulates the expression of an array of ROS-scavenging genes, whose feedback regulates its own levels in the nucleus and promotes OsNLP4-related functions. Our findings unveiled the novel central role of OsNLP4 linking the N-Fe balance and SL signaling, which fully demonstrates the ability of OsNLP4 to integrate and optimize nutrient signals to promote tillering to maximize grain yield and NUE in rice. Our findings provide the molecular mechanism for innovative fertilization with reduced N fertilizer input and significant returns in yield and environmental friendliness, which should have profound impacts on sustainable agriculture.

## Results

### OsNLP4 is a positive regulator of rice tillering

Our previous study identified OsNLP4 as a key player in rice yield and NUE (Wu *et al*., 2021). Field tests showed that *OsNLP4*-overexpressing plants (OE-1 and OE-9) exhibited more effective tillers under all different N conditions than the wild type (WT), whereas knockout (ko) mutants (*nlp4-1* and *nlp4-2*) had fewer tillers (Fig. 1a).

**Fig. 1.**
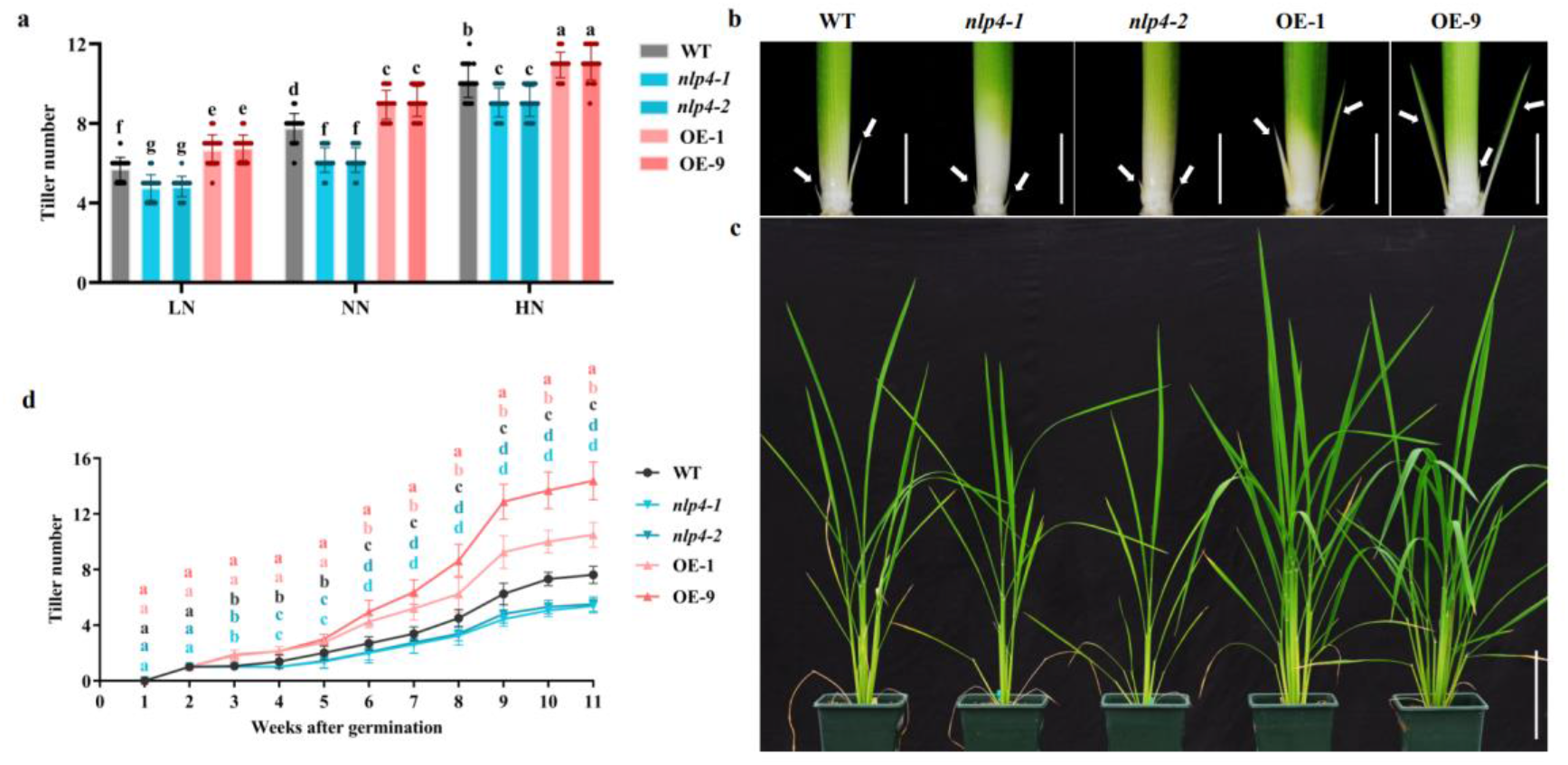
OsNLP4 positively regulates tillering in rice. **a**. Effective tiller numbers of WT, ko (*nlp4-1, nlp4-2*) and OE (OE-1, OE-9) plants in the field trial. The field trial was conducted under different N levels (LN: low N level with 80 kg urea per ha applied, NN: normal N level with 200 kg urea per ha, HN: high N level with 500 kg urea per ha). **b**. Tiller bud phenotypes of 2-week-old WT, ko and OE seedlings. Scale bar = 1 cm. **c**. Tiller phenotypes of 2-month-old WT, ko and OE plants grown in soil pots. Scale bar=15 cm. **d**. Tiller number curves of WT, ko and OE plants. The number of tillers were recorded once a week for 11 weeks. Values are the mean ± SD (n = 4 replicates and 18 plants per replicate for a, n = 16 plants for d). Different letters above bars denote significant differences (P < 0.05) from Duncan’s multiple range tests.

To confirm this result, we observed the formation of tiller buds in the WT, ko mutants (*nlp4-1, nlp4-2*) and OE (OE-1, OE-9) plants grown in soil pots and found that the OE plants formed tiller buds at a higher rate than WT and continued until 9 weeks after germination, whereas ko mutants formed fewer tillers than WT (Fig. 1b-d). By 11 weeks after germination, tiller number increased markedly in the OE plants by 37.8% (OE-1) and 88.5% (OE-9) compared with the WT but significantly decreased in the ko mutants by 29.5% (*nlp4-1*) and 27.9% (*nlp4-2*) (Fig. 1d). Together, these data unequivocally show that OsNLP4 is a positive regulator of rice tillering.

### OsNLP4 represses *OsD3* expression and downregulates SL signaling to promote tillering

To explore the mechanism by which OsNLP4 regulates tillering, we analyzed our RNA sequencing data and found that *OsD3* was a candidate target of OsNLP4. The transcript level of *OsD3* in the basal tissue where axillary buds emerge was elevated in ko mutants (*nlp4-1, nlp4-2*) compared with the WT, whereas it was decreased in the OE lines (Fig. 2a), suggesting that OsNLP4 represses *OsD3* expression. To confirm this, we conducted a series of assays. Chromatin immunoprecipitation (ChIP) assays showed that OsNLP4 directly bound to the P1 fragment containing the nitrogen-responsive element (NRE) in the *OsD3* promoter (Fig. 2b), which was further confirmed by an electrophoretic mobility shift assay (EMSA) (Fig. 2c). Dual-luciferase reporter experiments also showed that OsNLP4 repressed the transcription of *OsD3* (Fig. 2d). These results demonstrate that OsNLP4 represses *OsD3* expression and may consequently downregulate SL signaling. To confirm this, we measured the level of OsD53 targeted by OsD3 and the transcript levels of downstream *OsCKX9* and *OsFC1* in the basal tissue and found that the OsD53 levels significantly decreased in the ko mutants but obviously increased in the OE lines compared with the WT (Fig. 2e). The transcript levels of *OsCKX9* and *OsFC1* were upregulated in the ko mutants and downregulated in the OE lines, exhibiting a pattern similar to that of *OsD3* (Fig. 2f, g). These results indicate that SL signaling is downregulated in OE basal tissue.

**Fig. 2.**
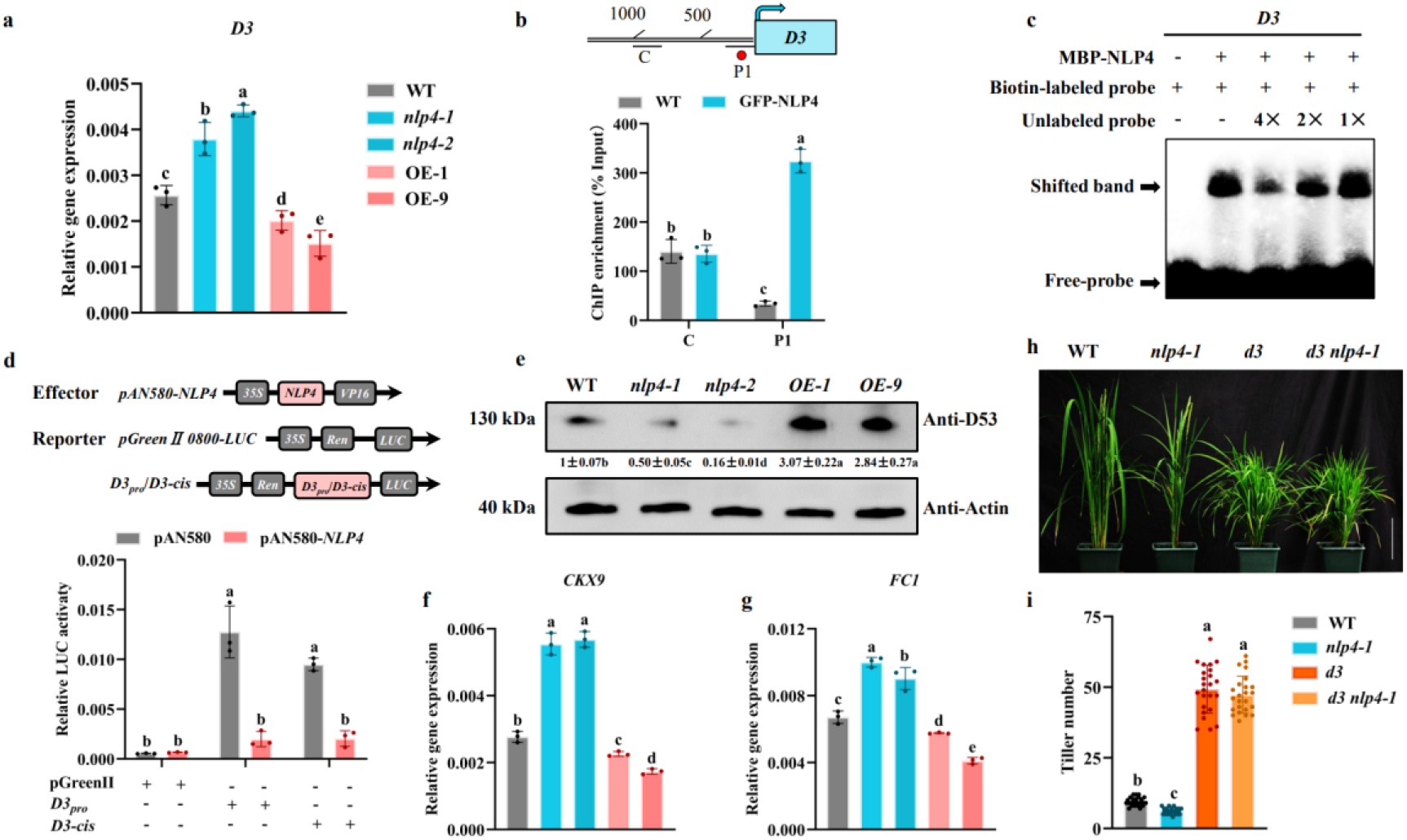
OsNLP4 promotes rice tillering by repressing *OsD3* in SL signaling. **a**. The expression levels of *OsD3* in basal tissues of WT, ko and OE plants cultured in soil for 30 days. **b**. The DNA binding rate of OsNLP4 to the *OsD3* gene promoter was analyzed by ChIP‒qPCR. **c**. EMSA results using a biotin-labeled NRE-like *cis*-fragment of the *OsD3* gene as a probe and an unlabeled fragment as a competitor. **d**. Dual-luciferase reporting transient assays showed that OsNLP4 inhibited the expression of *OsD3*. **e**. Protein levels of OsD53 in basal tissues of WT, ko and OE plants grown in soil detected by immunoblotting. The OsD53 protein level was quantified relative to the actin protein level by ImageJ. **f-g**. The expression levels of *OsCKX9* (f) and *OsFC1* (g) in basal tissues of WT, ko and OE plants. **h**. Tiller phenotypes of 2-month-old WT, *nlp4-1, d3* and *d3 nlp4-1* plants grown in soil pots. Scale bar=15 cm. **i**. Tiller numbers of plants as in h. Values are the mean ± SD (n = 3 replicates for a, b, d-g and n = 24 plants for i). Different letters above bars denote significant differences (P < 0.05) from Duncan’s multiple range tests.

To genetically confirm the negative regulation of *OsD3* by *OsNLP4*, we conducted genetic analysis of the double mutant *d3 nlp4-1*, which was created by editing *OsD3* in the *nlp4-1* background (Fig. S1). The double mutant exhibited a tillering phenotype similar to that of *d3* (Fig. 2h, i), indicating that *OsNLP4* and *OsD3* are in the same genetic pathway and that *OsNLP4* is upstream of *OsD3*. Taken together, these data clearly demonstrate that OsNLP4 positively regulates rice tillering by repressing *OsD3* in SL signaling.

### N-Fe balance modulates cellular H_2_O_2_ levels and OsNLP4 nuclear accumulation

Having confirmed that OsNLP4 is a positive regulator of rice tillering, we then addressed how the N-Fe balance modulates tillering. Our previous work has shown that OsNLP4 is N-responsive and coordinately regulates N and Fe metabolism pathways (our companion paper). If the N-Fe balance can modulate OsNLP4 levels in the nucleus, all growth and tiller phenotypes in response to N-Fe nutrition can be perfectly explained. Therefore, we examined OsNLP4 levels in the nucleus under different N-Fe conditions by observing the GFP signals in the *OsACTIN1pro:OsNLP4-GFP* rice roots. The GFP signal was hardly visible under LN-LFe conditions (Fig. 3a), whereas under HN-LFe conditions, the GFP signal was enhanced but remained invisible in the nucleus (Fig. 3b), indicating that the nitrate-induced nuclear localization of OsNLP4 requires the presence of Fe. This point was further supported by the dramatically increased nuclear GFP signals under HN-HFe conditions compared with those under HN-LFe conditions (Fig. 3b and f).

**Fig. 3.**
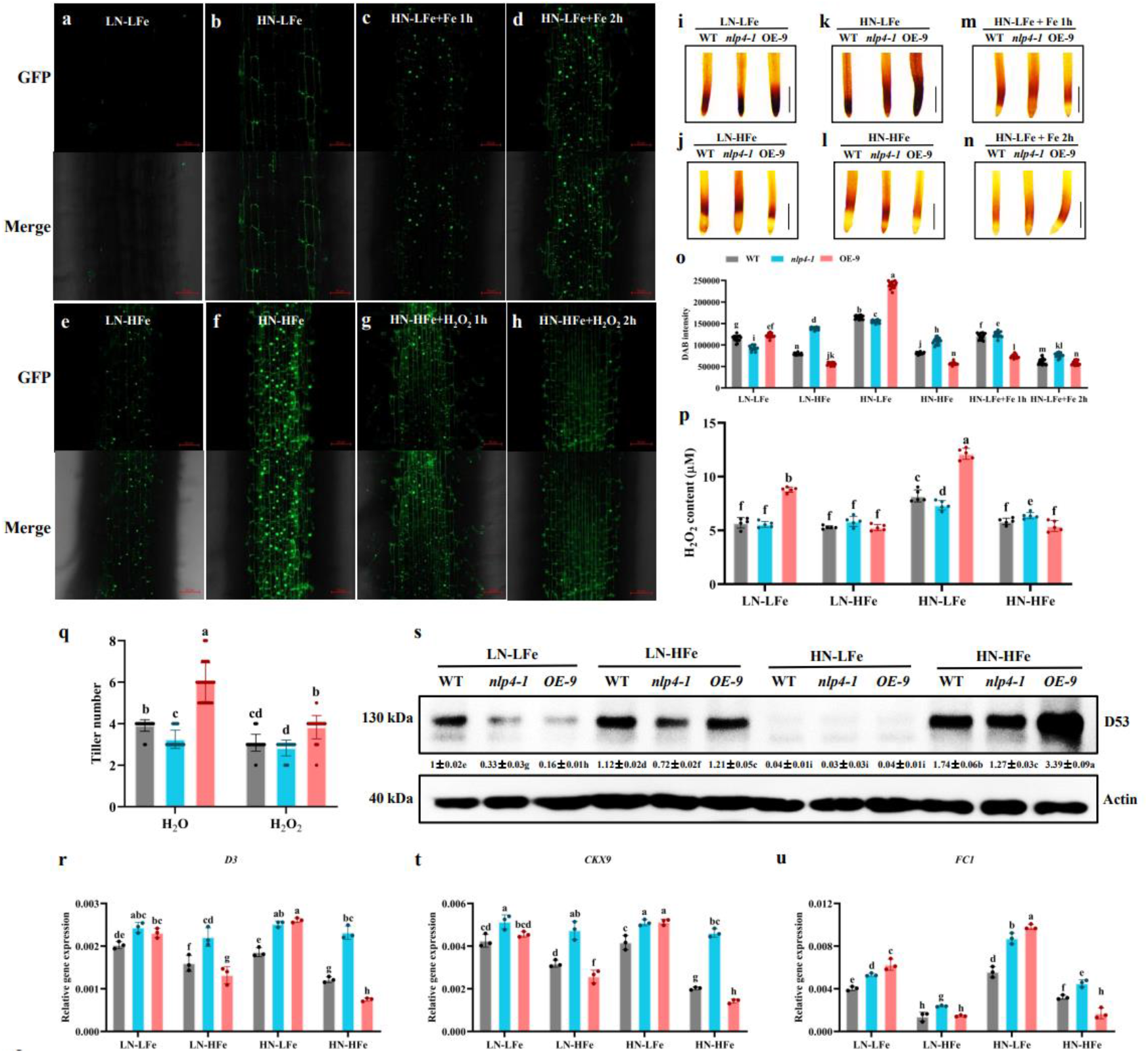
N-Fe balance modulates the H_2_O_2_ level and nuclear accumulation of OsNLP4. **a-h**. Subcellular localization of OsNLP4-GFP under LN-LFe (a), HN-LFe (b), HN-LFe + Fe 1 h (c) or 2 h (d), LN-HFe (e) and HN-HFe (f), HN-HFe + H_2_O_2_ 1 h (g) or 2 h (h) conditions. Scale bar = 50 μm. **i-n**. DAB staining of rice root tips grown under LN-LFe (i), LN-HFe (j), HN-LFe (k) and HN-HFe (l), HN-LFe + Fe 1 h (m) or 2 h (n) treatments. Scale bar = 1 cm. **o**. Quantification of DAB intensities in the images of i-n by ImageJ. **p**. The H_2_O_2_ concentration in the rice basal tissue. **q**. Exogenous supply of H_2_O_2_ inhibited rice tillering. **r-u**. The expression levels of *OsD3* (r), protein levels of OsD53 (s), and expression levels of *OsCKX9* (t) and *OsFC1* (u) in basal tissues of plants under different N-Fe conditions. The OsD53 protein level was quantified relative to the actin protein level by ImageJ. Values are the mean ± SD (n = 18 images for o, n = 5 replicates and 8 plants per replicate for p, n = 24 plants for q, and n = 3 replicates for r-u). Different letters above bars denote significant differences (P < 0.05) from Duncan’s multiple range tests.

To confirm these results, we supplied Fe to the plants grown under HN-LFe conditions and found that the GFP signal started to increase in the nucleus 1 hour after Fe was supplied (Fig. 3c) and was further enhanced by 2 hours (Fig. 3d). Moreover, under LN-HFe conditions, the GFP signals were much stronger and clearly detected in the nucleus than under LN-LFe conditions (Fig. 3a and e). All these results clearly demonstrate that Fe promotes OsNLP4 nuclear accumulation, and this promotion is enhanced when N-Fe is balanced.

H_2_O_2_ is negatively correlated with AtNLP6 and AtNLP7 nuclear accumulation (Chu et al., 2021). We treated transgenic plants grown under HN-HFe conditions with H_2_O_2_ and found that the GFP signal gradually diminished in the nucleus with time, indicating that H_2_O_2_ decreased OsNLP4 nuclear accumulation (Fig. 3g, h).

Taken together, these data show that balanced N-Fe conditions promote the nuclear accumulation of OsNLP4, whereas H_2_O_2_ reduces OsNLP4 levels in the nucleus, implying that N-Fe balance might affect H_2_O_2_ levels to modulate OsNLP4 levels in the nucleus and thereby its downstream events.

To prove this, we examined the H_2_O_2_ levels in the roots of different *OsNLP4* genotypes under different N-Fe conditions by 3,3’-diaminobenzidine tetrahydrochloride (DAB) staining (Fig. 3i-l). Under LN-LFe conditions, H_2_O_2_ accumulation exhibited fewer differences between all the genotypes compared with other N-Fe conditions (Fig. 3i, o). When the N-Fe was severely unbalanced (HN-LFe), H_2_O_2_ levels were markedly increased in the root tips, especially in OE plants (Fig. 3k, o). However, Fe addition to HN-LFe effectively reduced the H_2_O_2_ level within 2 h (Fig. 3m, n, o). Under HFe conditions, especially HN-HFe conditions, the H_2_O_2_ level was significantly lower than that under LFe conditions, with the OE plants responding most strongly (Fig. 3j, l, o). These results are basically consistent with the GFP signal data (Fig. 3a-f), suggesting that the N-Fe balance alters OsNLP4 levels in the nucleus by modulating H_2_O_2_ levels.

To confirm the above findings *in planta*, we measured the amount of H_2_O_2_ in rice basal tissue of plants with different *OsNLP4* genotypes under four N-Fe conditions and found that high endogenous H_2_O_2_ levels were associated with LFe conditions regardless of the N conditions, especially in the OE plants (Fig. 3p), consistent with the GFP signals (Fig. 3a-d). Furthermore, we performed exogenous H_2_O_2_ treatment in the early stage of rice tillering. The results showed that exogenously applied H_2_O_2_ could significantly inhibit rice tillering. The tiller number of the WT and OE plants decreased by 21% and 36%, respectively (Fig. 3q). Together, these results further confirm that the H_2_O_2_ level is negatively correlated with rice tillers and that HFe decreases the H_2_O_2_ level *in planta*.

We also investigated OsNLP4-regulated downstream events and found that the expression of *OsD3* was mostly repressed in the OE plants under HN-HFe conditions, while it was high in the OE lines under HN-LFe conditions (Fig. 3r). In contrast, in the *nlp4-1* plants, *OsD3* expression remained high under all N-Fe conditions. As the target protein of OsD3, the abundance of OsD53 was the highest when N-Fe was balanced and sufficient (HN-HFe), especially in the OE plants, and was the lowest under the poorest N-Fe balance (HN-LFe) (Fig. 3s). Consequently, the expression of *OsCKX9* and *OsFC1*, the downstream targets negatively regulated by OsD53, exhibited a very similar expression pattern to that of *OsD3* in the four N-Fe conditions (Fig. 3t, u).

Taken together, these data suggest that the N-Fe balance modulates OsNLP4 levels in the nucleus via H_2_O_2_, and OsNLP4 represses *OsD3* expression and downregulates SL signaling to promote tillering.

### OsNLP4 upregulates an array of genes involved in scavenging ROS

We conducted an in-depth analysis of RNA sequencing data and found that differentially expressed genes associated with H_2_O_2_ scavenging in the *OsNLP4* mutants and overexpressing lines were substantially enriched, including ascorbate peroxidases (*OsAPX1, OsAPX3, OsAPX5, OsAPX7*), catalases (*OsCATA, OsCATB*), glutathione peroxidase (*OsGPX1*), peroxidases (*OsPEROX4, OsPOX1*, etc.), and superoxide dismutase genes (*OsALM1, OsSOD*) (Fig. 4a). It is evident from the heatmap that these genes affecting the dynamics of H_2_O_2_ content exhibited similar changes under different N-Fe conditions, with the highest levels under HN-HFe conditions and the lowest levels under HN-LFe conditions, especially in the OE plants (Fig 4a). These results are consistent with the endogenous H_2_O_2_ levels that are inversely correlated with the expression levels of these genes (Fig. 3i-o) and that multiple NREs are located in the promoters of these genes (Table S1), suggesting that OsNLP4 may directly regulate the expression of these genes to affect cellular H_2_O_2_ levels. To confirm this, we chose *OsAPX7* as an example for further analysis. The expression level of *OsAPX7* was significantly upregulated in the OE plants and downregulated in the *nlp4-1* mutants (Fig. 4b). ChIP assays showed that the P1 fragment of the *OsAPX7* promoter was significantly enriched, confirming that OsNLP4 binds to the *OsAPX7* promoter *in vivo* (Fig. 4c). The binding was further confirmed by EMSA (Fig 4d). Subsequent dual-luciferase reporter assays showed that OsNLP4 activated the expression of the luciferase gene linked to the promoter of *OsAPX7* (Fig. 4e). Taken together, these results suggest that OsNLP4 can upregulate the genes involved in scavenging H_2_O_2_, such as *OsAPX7*, to counteract H_2_O_2_ according to the N-Fe conditions, providing a feedback loop to increase OsNLP4 levels in the nucleus.

**Fig. 4.**
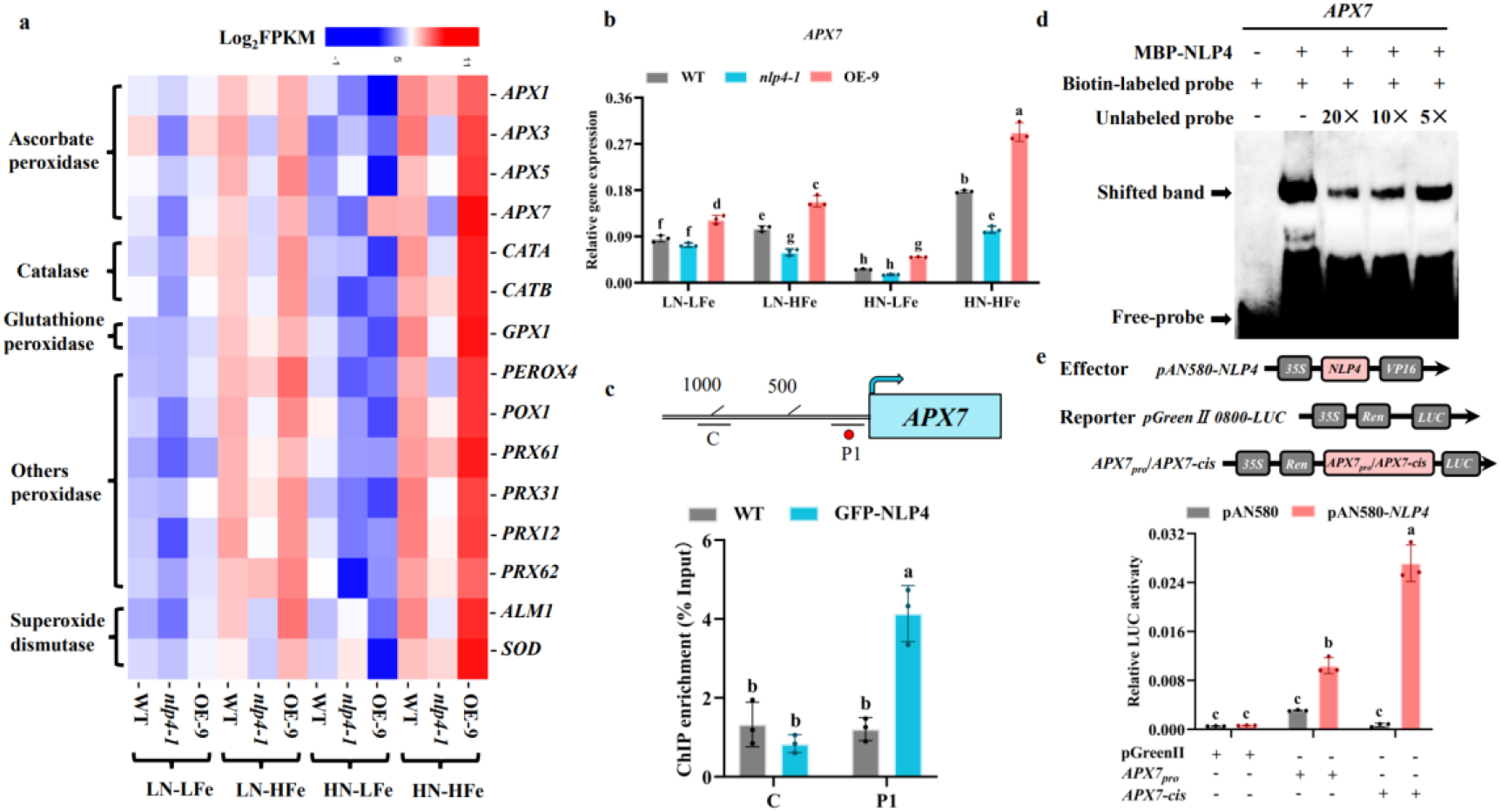
OsNLP4 upregulates an array of genes involved in scavenging H_2_O_2_. **a**. Hierarchical cluster analysis was performed on the expression data of 15 genes related to H_2_O_2_ in the WT, ko (*nlp4-1*), and OE (OE-9) under different N-Fe conditions. The value of the blue to red gradient bar represents the log2 of the ratio. Only results with a P value < 0.05 that were confirmed in three independent experiments were included. A color code was used to visualize the data. FPKM: fragments per kilobase of transcript per million fragments mapped. **b**. The relative expression levels of the *OsAPX7* gene in 12-day-old WT, ko and OE plants (basal tissue) under different N-Fe conditions. **c**. The DNA binding rate of OsNLP4 to the *OsAPX7* gene promoter was analyzed by ChIP‒qPCR. **d**. EMSA results using a biotin-labeled NRE-like cis-fragment of the *OsAPX7* gene as a probe and an unlabeled fragment as a competitor. **e**. Dual-luciferase reporting transient assays showed that OsNLP4 induced the expression of *OsAPX7*. Values are the mean ± SD (n = 3 replicates for c and e). Different letters above bars denote significant differences (P < 0.05) from Duncan’s multiple range tests.

## Discussion

Tillering has attracted much attention because of its direct contribution to grain yield (Wang *et al*., 2022). Among the many factors that affect tillering, nutrients and hormones have always been a research hotspot. In this study, we revealed that OsNLP4 is a positive regulator of rice tillering (Fig. 1, 2). Balanced N-Fe strongly induced the nuclear accumulation of OsNLP4 by reducing the level of H_2_O_2_ (Fig. 3), thereby inhibiting the expression of downstream *OsD3* and SL signaling and finally promoting rice tillering. In addition, OsNLP4 can regulate the expression of an array of ROS-scavenging genes, such as *OsAPX7*, to attenuate H_2_O_2_ levels (Fig. 4), providing a feedback loop that promotes its own nuclear accumulation. Our findings reveal a novel mechanism by which OsNLP4 integrates the N-Fe nutrient balance with SL signaling to promote rice tillering, thereby maximizing yield and NUE, which will exert profound impacts on the improvement of grain yield and NUE that benefit sustainable green agriculture.

### OsNLP4 promotes rice tillering by repressing *OsD3* expression

NLPs are pivotal transcription factors regulating N fixation and cold stress responses and act as nitrate sensors, as recently shown for AtNLP7 (Ding et al., 2022; Jiang et al., 2021; Liu et al., 2022). However, their role in tillering regulation has not been reported to date. Compared with WT, the *OsNLP4*-OE plants exhibited faster tiller occurrence and more tillers, while the ko mutants had fewer tillers (Fig. 1), indicating that OsNLP4 positively regulates tillering in rice.

SLs are important phytohormones that regulate tillering in rice (Wang *et al*., 2020). As one of the core components of SL signaling (Jiang *et al*., 2013; Zhou *et al*., 2013), *OsD3* expression is directly regulated by OsNLP4. Molecular and biochemical assays demonstrated that OsNLP4 transcriptionally represses *OsD3* expression by binding to an NRE element within its promoter; therefore, *OsD3* is significantly downregulated in the OE plants but upregulated in the ko mutants (Fig. 2a-d). OsD3 forms a complex with the SL receptor OsD14 to regulate the stability of the OsD53 protein, thus affecting the expression of downstream branching inhibitory genes (Jiang *et al*., 2013; Zhou *et al*., 2013). Compared with WT, the protein levels of OsD53 were significantly increased in the OE plants and reduced in the ko mutants, resulting in its target genes *OsCKX9* and *OsFC1* showing a similar expression pattern to *OsD3* (Duan *et al*., 2019; Fang *et al*., 2020), ultimately causing corresponding changes in tiller numbers (Figs. 1, 2a, e-g). The *d3 nlp4* double mutants exhibited a high-tillering phenotype similar to that of *d3* (Fig. 2h, i), providing genetic evidence that *OsNLP4* is epistatic to *OsD3*. Taken together, the OsNLP4-OsD3 module enables the level of OsNLP4 in the nucleus to control the intensity of the inhibition of *OsD3* expression, subsequently affecting tillering.

### N-Fe balance modulates cellular H_2_O_2_ levels that negatively affect OsNLP4 nuclear accumulation

The level of OsNLP4 in the nucleus is regulated by the N-Fe balance. We previously reported that the nuclear accumulation of OsNLP4 was positively correlated with the availability of nitrate under sufficient Fe (Wu *et al*., 2021). In this study, we found that the presence of Fe is required for nitrate-induced nuclear localization of OsNLP4. Under the HN-LFe conditions, OsNLP4 was almost absent in the nucleus, whereas the nuclear accumulation level of OsNLP4 was the highest under balanced N-Fe (HN-HFe) conditions (Fig. 3a, b, e, f). Fe supplementation to HN-LFe conditions clearly promotes the nuclear localization of OsNLP4 (Fig. 3c, d), again clarifying the necessity of the N-Fe balance for OsNLP4 nuclear accumulation.

Interestingly, we found that different N-Fe conditions were accompanied by varying H_2_O_2_ levels in rice roots and tiller bases, which were negatively correlated with OsNLP4 nuclear accumulation (Fig. 3). N deprivation is known to increase H_2_O_2_ in distinct areas of the plant, such as shoot and root apical meristems (Wany et al., 2018). In this study, we found that the Fe concentration is critical as well. When Fe was deficient, N sufficiency exacerbated the impact of Fe depletion and induced H_2_O_2_ bursts, especially in the OE plants (Fig. 3i-l, o, p).

Fe serves as the intrinsic constituent or metal cofactor in H_2_O_2_-scavenging enzymes such as catalase, phenolic-dependent peroxidases, ascorbate peroxidases and Fe superoxide dismutase (Briat *et al*., 2015). Fe deficiency leads to a decrease in the activities of these enzymes, as shown for peroxidase activity (Ranieri et al., 2001). In addition, the Fenton reaction of Fe with H_2_O_2_ is certainly an important direct contributor to the reduction of H_2_O_2_ (Jeong and Guerinot, 2009). Consistent with this, LFe was always associated with high H_2_O_2_ content regardless of N concentration (Fig. 3i-l, o, p), whereas HFe reduced H_2_O_2_ levels *in planta*, as confirmed by DAB staining after Fe supplementation of HN-LFe-treated rice (Fig. 3m, n, o). These data support that the N-Fe balance significantly modulates H_2_O_2_ levels *in planta*.

H_2_O_2_ suppresses nitrate signaling by modulating the nucleocytoplasmic shuttling of AtNLP7, a homolog of OsNLP4 in *Arabidopsis* (Chu *et al*., 2021). The nuclear fluorescence signal of OsNLP4-GFP was significantly reduced within 2 h of H_2_O_2_ treatment, indicating that H_2_O_2_ could inhibit the nuclear accumulation of OsNLP4 (Fig. 3f-h). Therefore, the variation in H_2_O_2_ content under different N-Fe conditions regulated the level of OsNLP4 in the nucleus, resulting in all *OsNLP4* genotypes showing different *OsD3* expression patterns and downstream events, accompanied by corresponding changes in tiller number, which is consistent with the yield and NUE results in our companion paper (Fig. 3r-u, Fig. 2 of companion paper).

H_2_O_2_ posttranslationally modifies target proteins by oxidizing the thiol groups of cysteine (Cys) residues, thereby changing their structural configuration and function (Tian et al., 2018). The OsNLP4 protein contains 20 Cys residues, and Cys213 and Cys236 are located in the N-terminal GAF-like domain critical for nitrate sensing (Liu *et al*., 2022; Wu *et al*., 2021). Oxidative modification of these Cys residues by H_2_O_2_ may interfere with phosphorylation of serine sites of this domain by calcium-dependent protein kinases (CPKs), inhibiting the nuclear localization of OsNLP4 (Ding *et al*., 2022; Liu et al., 2017). In addition, Cys801 and Cys808 are located in the PB1 domain responsible for protein‒protein interactions, and their oxidation may hinder the interaction of partners responsible for helping OsNLP4 nuclear accumulation, such as OsSPX4 (Hu *et al*., 2019). Alternatively, H_2_O_2_ may directly oxidize these partners and thus indirectly inhibit OsNLP4 nucleocytoplasmic shuttling.

In addition, H_2_O_2_ inhibited the nuclear accumulation of OsNLP4, which in turn regulated the expression of multiple ROS-scavenging genes affecting H_2_O_2_ levels, such as *OsAPX7*, thus forming a feedback regulatory loop and amplifying the cascade signal of the N-Fe balance-H_2_O_2_-OsNLP4-OsD3 module (Fig. 4). Taken together, balanced N-Fe promotes the nuclear accumulation of OsNLP4 by reducing cellular H_2_O_2_ levels, thereby repressing *OsD3* expression and downstream SL signaling and ultimately increasing rice tillers.

In conclusion, we have uncovered a novel mechanism by which OsNLP4 integrates the N-Fe balance and phytohormone SL signaling to regulate tillering in rice. A working model is proposed to illustrate the central role of OsNLP4 in the regulation of rice tillering in response to the N-Fe nutrient balance (Fig. 5). Our findings should have significant and profound impacts on the improvement of crop yield and NUE by manipulating NLPs and innovative fertilization formulas and methods with reduced N fertilizer input but boosted grain yield, which will greatly benefit green and sustainable agriculture worldwide.

**Fig. 5.**
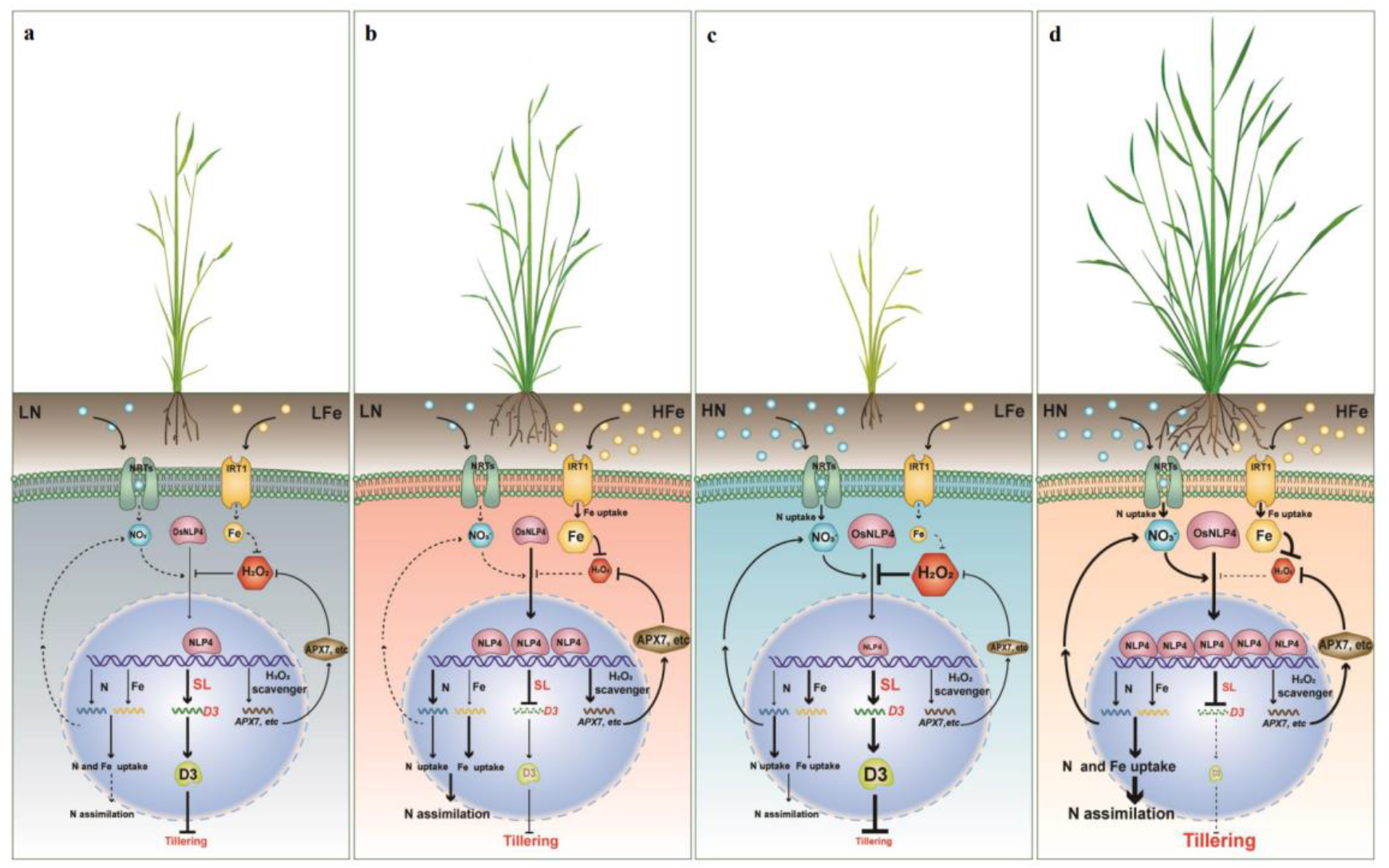
A working model - OsNLP4 integrates N-Fe nutrient signals with SL signaling to regulate rice tillering. **a**. Under LN-LFe conditions, LFe-induced H_2_O_2_ accumulation together with N starvation inhibits the nuclear localization of OsNLP4, failing to regulate downstream target genes. The inhibition of the tillering-suppressive gene *OsD3* is relieved, while the absorption and utilization of N-Fe and the scavenging ability of H_2_O_2_ are decreased, which severely impairs the growth of rice, with leaf chlorosis and fewer tillers. **b**. Under LN-HFe conditions, the reduction of H_2_O_2_ by HFe partially alleviates the inhibition of nuclear accumulation of OsNLP4 by LN, which therefore slightly enhances the inhibition of *OsD3* while upregulating *OsAPX7* to promote H_2_O_2_ scavenging, resulting in increased tillering to some extent compared to the LN-LFe conditions. **c**. Under HN-LFe conditions, the N-Fe imbalance is exacerbated by the contradiction between the urgent demand of Fe for N metabolism and poor Fe availability, producing the highest H_2_O_2_ levels among the four N-Fe conditions, which strongly inhibits the nuclear accumulation of OsNLP4, resulting in the lowest levels of OsNLP4 in the nucleus even under HN, thereby turning off all downstream events, ultimately leading to a sharp reduction in tiller number and the most severe leaf chlorosis. **d**. Under HN-HFe conditions, HN promotes the nuclear accumulation of OsNLP4, while HFe inhibits the level of H_2_O_2_, resulting in the highest level of OsNLP4 in the nucleus, strengthening the regulation of target genes involved in the N, Fe, H_2_O_2_ and SL signaling pathways, which promotes the coordinated utilization of N and Fe nutrients, inhibits *OsD3* expression and downstream SL signaling, and maintains the low level of H_2_O_2_ by upregulating the expression of ROS-scavenging genes such as *OsAPX7*, ultimately greatly maximizing rice tillering and growth. Black arrows indicate positive effects. Bar-headed lines represent negative effects. Dashed lines represent weak routes of corresponding solid lines and arrows. The thickness of the lines and the size of the font represent the degree of positive or negative effects. The size of the shapes represents the levels.

## Methods

### Plant materials

All rice (*Oryza sativa*) plants used in this study were derived from the japonica variety Zhonghua 11 (ZH11). *OsNLP4* ko mutants and overexpressing plants were previously used lines (Wu *et al*., 2021). The *OsACTIN1:OsNLP4* overexpression construct inserts the *OsNLP4* coding region into pCB2006 through the GATEWAY clone system. The homozygous OE-1 and OE-9 overexpression lines (T4 generation) were obtained by *Agrobacterium tumefaciens* (EHA105) transformation and glufosinate screening. The mutants *nlp4-1, nlp4-2, d3* and *d3 nlp4-1* were obtained from Bogle Hangzhou Co., Ltd. (Hangzhou, China) (http://www.biogle.cn/) using CRISPR/Cas9 technology.

### Plant growth conditions

#### Seedlings in soil culture

Seeds of all plants (WT, *nlp4-1, nlp4-2*, OE-1, OE-9, *d3* and *d3*/*nlp4-1*) were washed with distilled water and incubated at 37 °C for 3 days. Germinated seeds were transferred to pots filled with soil (the pot dimensions were 15×15×15 cm^3^). Each treatment contained 50 pots for each genotype. A single plant was grown per pot. The provided growth conditions were kept at a 12-h light (30 °C)/12-h dark (28 °C) cycle in a greenhouse.

#### Seedlings in hydroponic culture

Seeds of WT, *nlp4-1*, and OE-9 were washed with distilled water and incubated at 37 °C for 3 days. Germinated seeds were transferred to modified Kimura B solution (pH 5.8) with different N and Fe availability treatments (LN: 0.05 mM KNO_3_, HN: 5 mM KNO_3_, LFe: 1 μM Fe(II)- EDTA, HFe: 100 μM Fe(II)-EDTA) to grow for 1-2 weeks. Nutrient solutions were replaced every two days. The provided growth conditions were kept at a 16-h light (30 °C)/8-h dark (28 °C) cycle in a climate chamber.

#### Seedlings in vermiculite for long-term treatment

Seeds of WT, *nlp4-1*, and OE-9 were washed with distilled water and incubated at 37 °C for 3 days. Germinated seeds were transferred to pots filled with vermiculite (the pot dimensions were 15×15×15 cm^3^). A single plant was grown per pot. Each treatment contained 50 pots for each genotype. Plants were fed different N-Fe solutions (LN: 1 mM KNO_3_, HN: 5 mM KNO_3_, LFe: 50 μM Fe(II)-EDTA, HFe: 200 μM Fe(II)-EDTA) until maturity. Each treatment was irrigated with 10 L of nutrient solution each time. During the whole growth period, plants were irrigated 12 times at intervals of 12 days. The provided growth conditions were kept at a 12-h light (30 °C)/12-h dark (28 °C) cycle in a greenhouse.

### Field tests of rice

Rice field experiments were conducted in Chengdu (Sichuan Province, China) in 2019 in paddy fields under natural growing conditions. The field experiment comprised five OsNLP4 genotypes (wild type, ko mutant *nlp4-1*/*nlp4-2*, and overexpression line OE-1/OE-9) and three N treatments: urea was used as the N source with 80 kg/ha for low N (LN), 200 kg/ha for normal N (NN) and 500 kg/ha for high N (HN). The plants were transplanted into 200 plants for each plot (8 m^2^), and four replicates were used for each N condition. To reduce the variability in the field test, the fertilizers were evenly applied to every plot for each N application level. For data collection, the edge lines of each plot were excluded to avoid margin effects.

### RNA-seq

Each strain (approximately 96 seedlings) with different N-Fe (0.05/5 mM KNO_3_ and 1/100 μM Fe(II)-EDTA) treatments was hydroponically grown in an artificial climate chamber under the conditions described above. Two-week-old seedlings (whole plants) were selected for RNA sequencing. For each treatment, 30 seedlings were collected as a sample, and three independent biological replicates were conducted. RNA library construction and sequence analysis were described previously (Wu *et al*., 2021).

### qRT‒PCR analysis

Total RNA was isolated from 1-month-old rice (basal tissue) grown in soil or different N-Fe hydroponic solutions in vermiculite (LN: 1 mM KNO_3_, HN: 5 mM KNO_3_, LFe: 50 μM Fe(II)-EDTA, HFe: 200 μM Fe(II)-EDTA) using TRIzol reagent (TRANSGEN, cat # ET111). Full-length cDNA was then reverse transcribed using a cDNA synthesis kit (TRANSGEN, cat # AE311-02). The qRT‒PCR step one Plus real-time PCR system was used for the TaKaRa SYBR premix Ex Taq II reagent kit (cat # Q111-02). Subsequent qRT‒PCR procedures were performed according to the manufacturer’s instructions, and each qRT‒PCR assay was repeated at least three times with three independent RNA preparations. Rice *Actin1* was used as an internal reference. The primers used are shown in Table S2.

### ChIP-PCR assay

Fourteen-day-old seedlings (whole plants) in HN-HFe hydroponic culture were taken for the ChIP experiment (HN: 5 mM KNO_3_ and HFe: 100 μM Fe(II)-EDTA). Samples (3-4 g) of *OsACTIN1pro:OsNLP4-GFP* transgenic plants (whole seedlings) were fixed and crosslinked with 1% formaldehyde vacuum for 10 min, and then 125 mM glycine was added to stop the reaction. The samples were frozen in liquid nitrogen and then ground into powder. Ultrasound at low temperature (4 °C) fragmented chromatin to an average size of 400-600 bp. GFP-trap magnetic beads are used for immunoprecipitation of protein DNA complexes. Chromatin without antibody precipitation was used as an internal reference, and the gene enrichment level was detected by real-time PCR. The sequences of related primers are shown in Table S2.

### Protoplast transfection and dual-luciferase reporter assay

The stems of rice grown in soil for 2-3 weeks were used to prepare protoplasts. In protoplast transient expression experiments, plasmids were transfected into protoplasts as described previously (Chu *et al*., 2021). One thousand bp upstream of the target gene initiation codon and the NRE-like element (*cis*-fragment) were cloned into the vector pGreenII 0800-LUC to generate a reporter gene for dual-luciferase detection. The full-length *OsNLP4* CDS is inserted into the pAN580 vector to generate the effector. Firefly luciferase (LUC) activity and Renilla luciferase (REN) activity were determined 12-18 h after transfection using a dual luciferase reporting system (Vazyme, cat # DL101-01). The LUC and REN activity ratios were determined at least three times.

### Electrophoretic mobility shift assay (EMSA)

EMSAs were performed as described (Liu *et al*., 2021). The *OsNLP4*-coding region was cloned into the pET30a vector, and MBP-OsNLP4 fusion protein was expressed in the *Escherichia coli* Rosseta strain. Purified recombinant MBP-OsNLP4 protein and MBP protein from *Escherichia coli. Cis*-fragments of target gene promoters were synthesized, and biotin was labeled at the DNA ends. Unlabeled fragments of biotin with the same sequence were used as competitors. EMSAs were performed with LightShift® Chemiluminescent EMSA Kits (cat # 20148) from Thermo Scientific company. Each reaction was loaded onto a 2% native agarose gel in 0.5 x TBE buffer for electrophoresis. The results were detected using a CCD camera system (GE Healthcare, Image Quant LAS 4000).

### DAB staining

Each strain (approximately 48 seedlings) with four N-Fe treatments (0.02/10 mM KNO_3_ and 1/200 μM Fe(II)-EDTA) was hydroponically grown in an artificial climate chamber under the conditions described above. The Fe concentration of the Fe-supplemented group (HN-LFe + Fe) was 200 μM, and the samples were stained after incubation on a shaker for 2 h. Roots of 1-week-old seedlings were incubated with 1 mg/mL DAB solution for 2 h at room temperature. Samples were then immersed in 80% (v/v) ethanol for 10 min to terminate the staining before being photographed. The signal intensity was quantified by ImageJ (Chu *et al*., 2021).

### H_2_O_2_ measurement

The H_2_O_2_ concentration in the tiller base of rice was measured by a Hydrogen Peroxide Assay Kit (Red Hydrogen Peroxide/Peroxidase Assay Kit, Amplex™, cat # A22188). The tiller bases of rice grown for four N-Fe treatments (1 mM/5 mM KNO_3_, 50/200 μM Fe(II)-EDTA) were used as experimental materials. The 1-month-old samples were frozen in liquid nitrogen and ground into powder. After being dissolved in PBS buffer, the samples were centrifuged at 12000 rpm at 4 °C for 10 min. The supernatant was collected, and the content was analyzed according to the manufacturer’s protocol. The OD value was measured at 560 nm wavelength by a multifunction microplate reader iD5.

### Subcellular localization of OsNLP4-GFP

To investigate the subcellular localization of OsNLP4, *OsACTIN1pro:OsNLP4-GFP* transgenic seedlings were germinated and grown in modified Kimura B medium with four N-Fe treatments (0.02/10 mM KNO_3_ and 1/200 μM Fe(II)-EDTA) for 14 days. Then, HN-LFe-fed rice was treated with 10 mM Fe(II)-EDTA for 2 h, and HN-HFe-fed rice was treated with 1 mM H_2_O_2_ for 2 h. Laser scanning confocal imaging was performed using a Chase 880 microscope equipped with an argon laser (488 nm for GFP excitation).

### Tillering assay of exogenous H_2_O_2_ treatment

Two-week-old nontillered rice plants grown under HN-HFe (HN: 5 mM KNO_3_, HFe: 200 μM Fe(II)-EDTA) conditions were used in this experiment. Cubic sponges of the same size (16 cm^3^) were squeezed out of the air and soaked in the prepared H_2_O_2_ solution (1 mM) until fully saturated for this experiment. The H_2_O_2_ solution was freshly prepared before use each time. Water-soaked sponges were used as controls. The soaked sponges were wrapped around the base of the rice plants with a wire and replaced once a day. After a week of treatment, the sponges were removed, and the tillers were counted.

### Western blotting assay

RIPA lysis buffer (strong) (Beyotime, China, cat # P0013B) was used to extract proteins from the tiller bases of rice (30 days) grown in soil or vermiculite pots under four N-Fe treatments (0.02 or 10 mM KNO_3_ and 1 or 200 μM Fe(II)-EDTA). The seedlings were frozen in liquid nitrogen and ground into powder. The protein extract was separated by SDS‒PAGE and transferred to a nitrocellulose membrane. The antibodies used in western blotting were as follows: anti-OsD53 (rabbit pAb, antibodies prepared by Xuelu Wang Laboratory, Henan University, Kaifeng, China) (Fang *et al*., 2020), 1:1000 for western blotting. Anti-ACTIN antibody (M20009, Mouse mAb, Abmart, Shanghai, China) at 1:1000 was used for western blotting. Goat anti-mouse lgG-HRP (M21001, Abmart, Shanghai, China) and goat anti-rabbit lgG-HRP (M21002, Abmart, Shanghai, China) were used at 1:5000 for western blotting. Image Quant LAS 4000 (GE, USA), as the CCD camera system, was used for band intensity quantification with the Super Signal West Femto Trial Kit (Thermo, Rockford, IL, USA, cat # 34095).

### Statistical analysis

The data were statistically analyzed, and multiple comparisons were performed using the Duncan multiple range test. A P value less than 0.05 was considered statistically significant.

## Data and materials availability

The data supporting the findings of this study are available within this paper and its Supplementary information files. The sequence data used in this study can be found in the Rice Genome Annotation Project (https://rice.plantbiology.msu.edu/) under the following accession numbers: *OsNLP4, LOC_Os09g37710*; *OsD3, LOC_Os06g06050*; *OsCKX9, LOC_Os05g31040*; *OsFC1, LOC_Os03g49880*; *OsAPX7, LOC_Os04g35520*. The datasets and materials used during the current study are available from the corresponding author on reasonable request.

## Author Contributions

J.W. and C.B.X. designed the experiments. Y.S., G.Y.W., J.X.W. and J.W. performed the experiments and data analyses. Y.S. wrote the manuscript. C.B.X., J.W., and L.H.Y. revised the manuscript, C.B.X. supervised the project.

## Acknowledgments

We thank Dr. Xuelu Wang (College of Life Sciences, Henan University, Kaifeng, Henan, China) for providing the antibodies against OsD53, Dr. Chengcai Chu (Institute of Genetics and Developmental Biology, CAS, Beijing, China) for his critical comments and suggestions on the manuscript, Drs. Peng Qin and Shigui Li for their assistance with field trials, and Dr. Chunming Wang (Nanjing Agricultural University, Nanjing, China) for providing the pAN580 plasmid.

## Supplementary figures and tables

**Figure S1.**
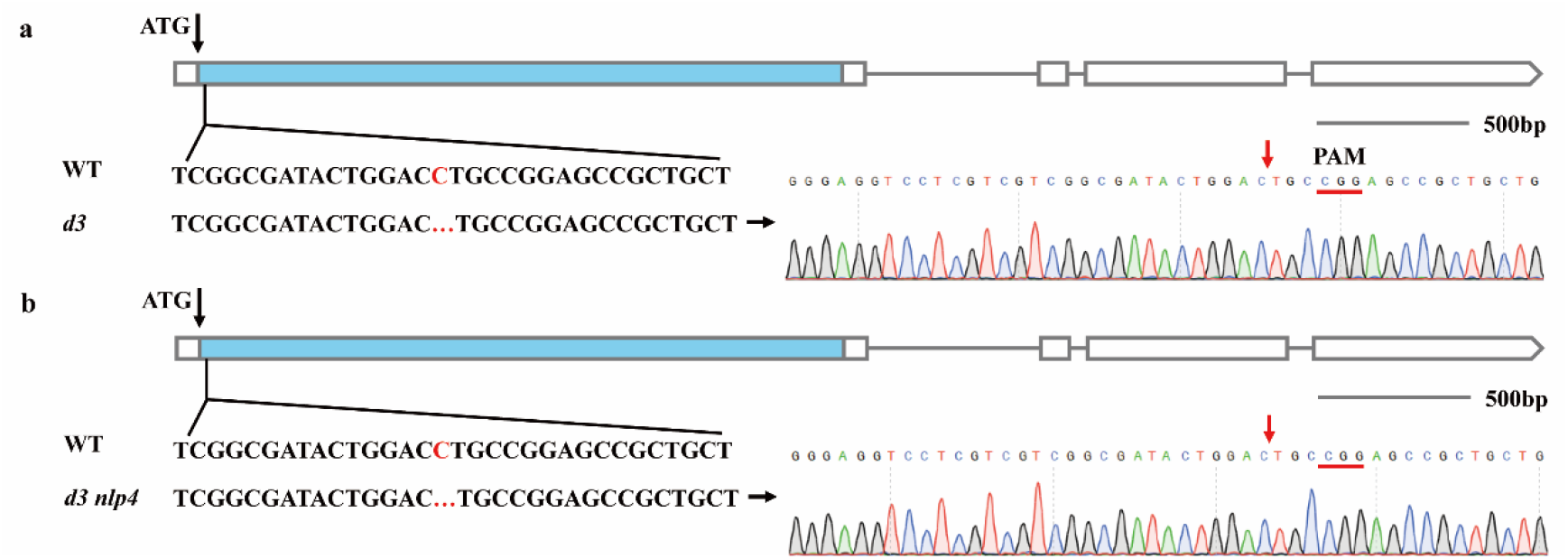
Verification of the CRISPR/Cas9-edited mutations in *OsD3* of *d3* and *d3 nlp4-1* mutants. **a-b**. Base changes or deletion of the knockout mutants generated by CRISPR‒Cas9-based editing were verified by sequencing *OsD3* cDNA from *d3* (a) and *d3 nlp4-1* mutants (b).

**Table S1.**
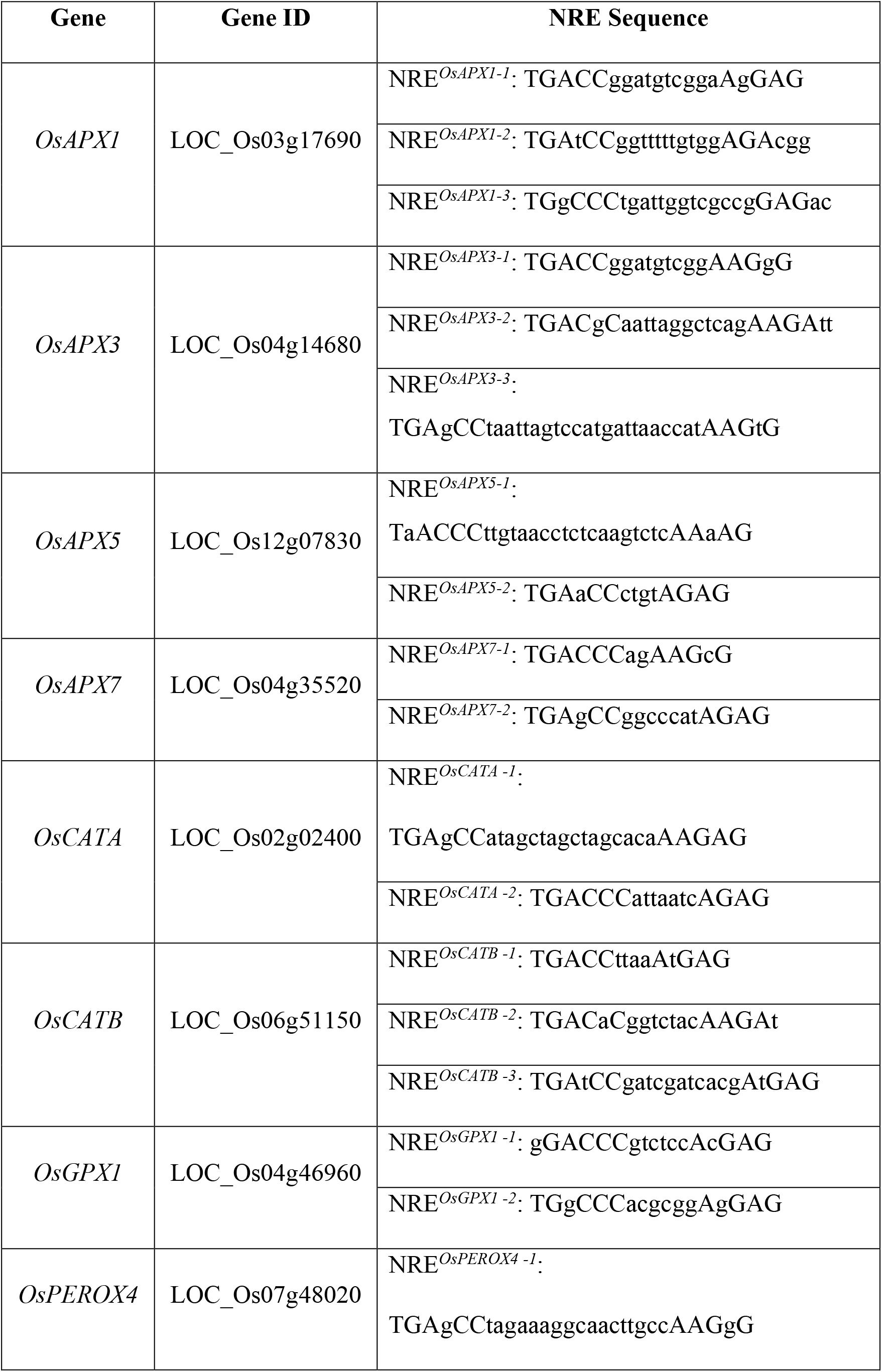
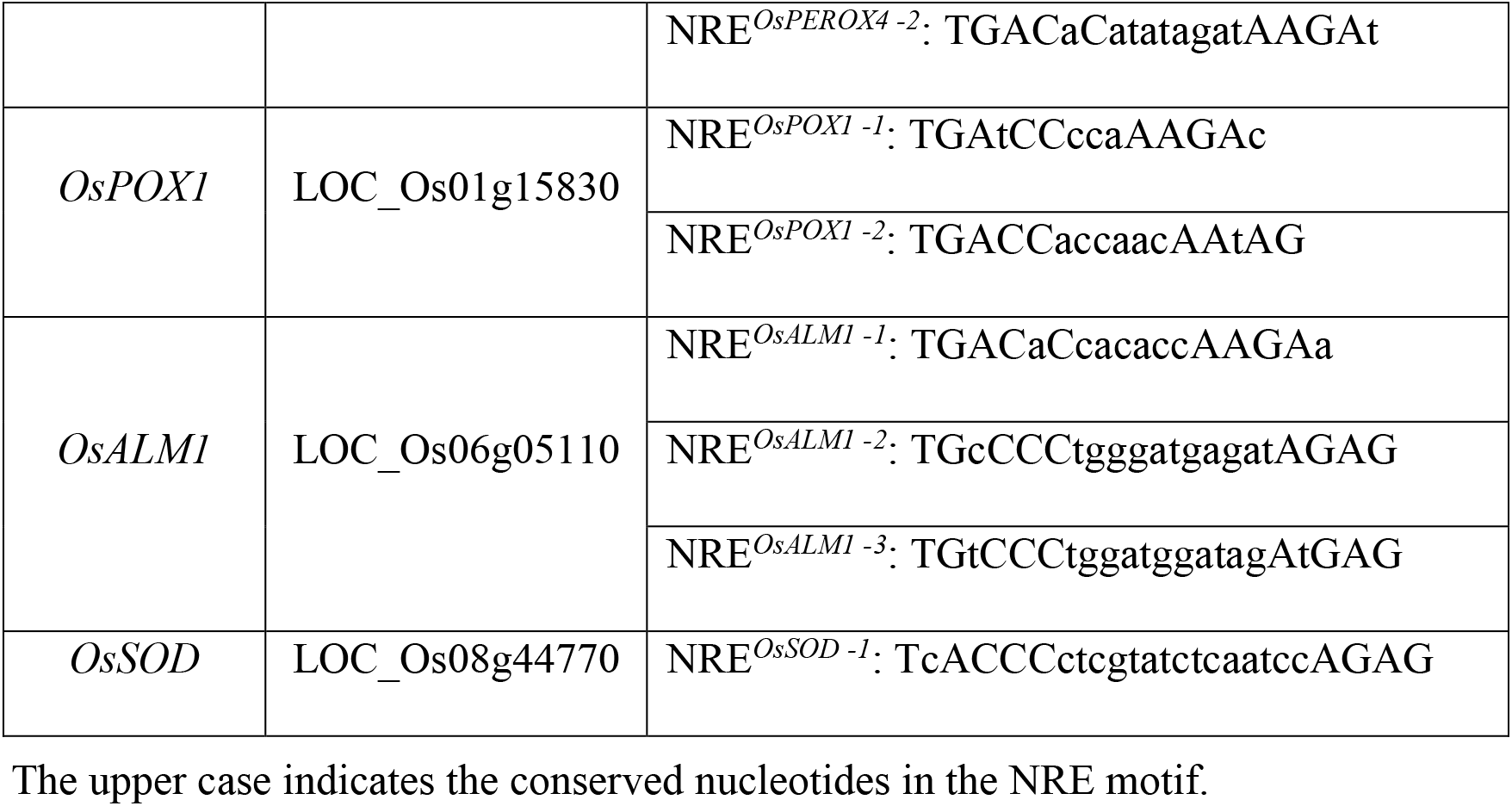
Predicted NREs in the promoter of H_2_O_2_-scavenging genes.

**Table S2.**
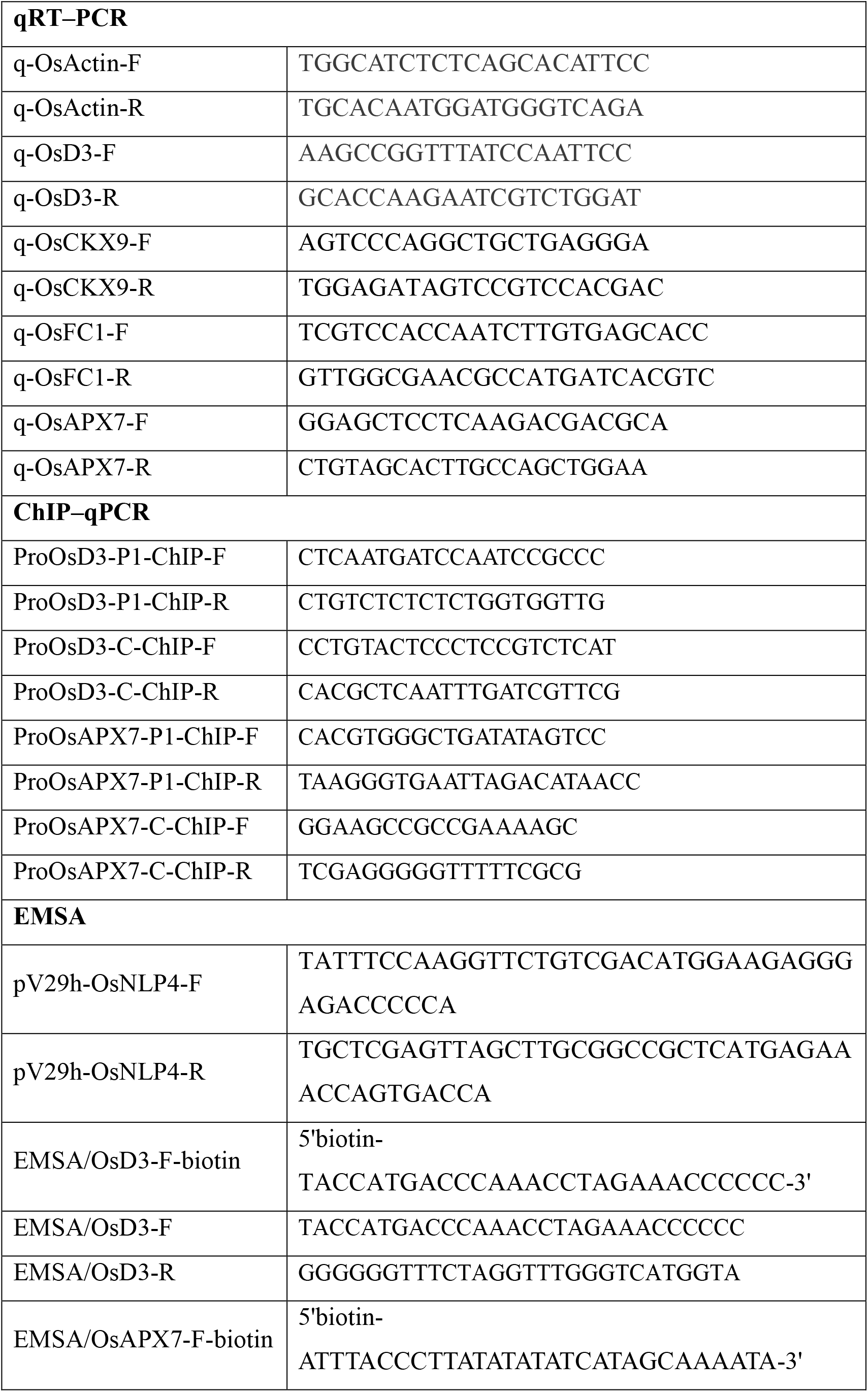
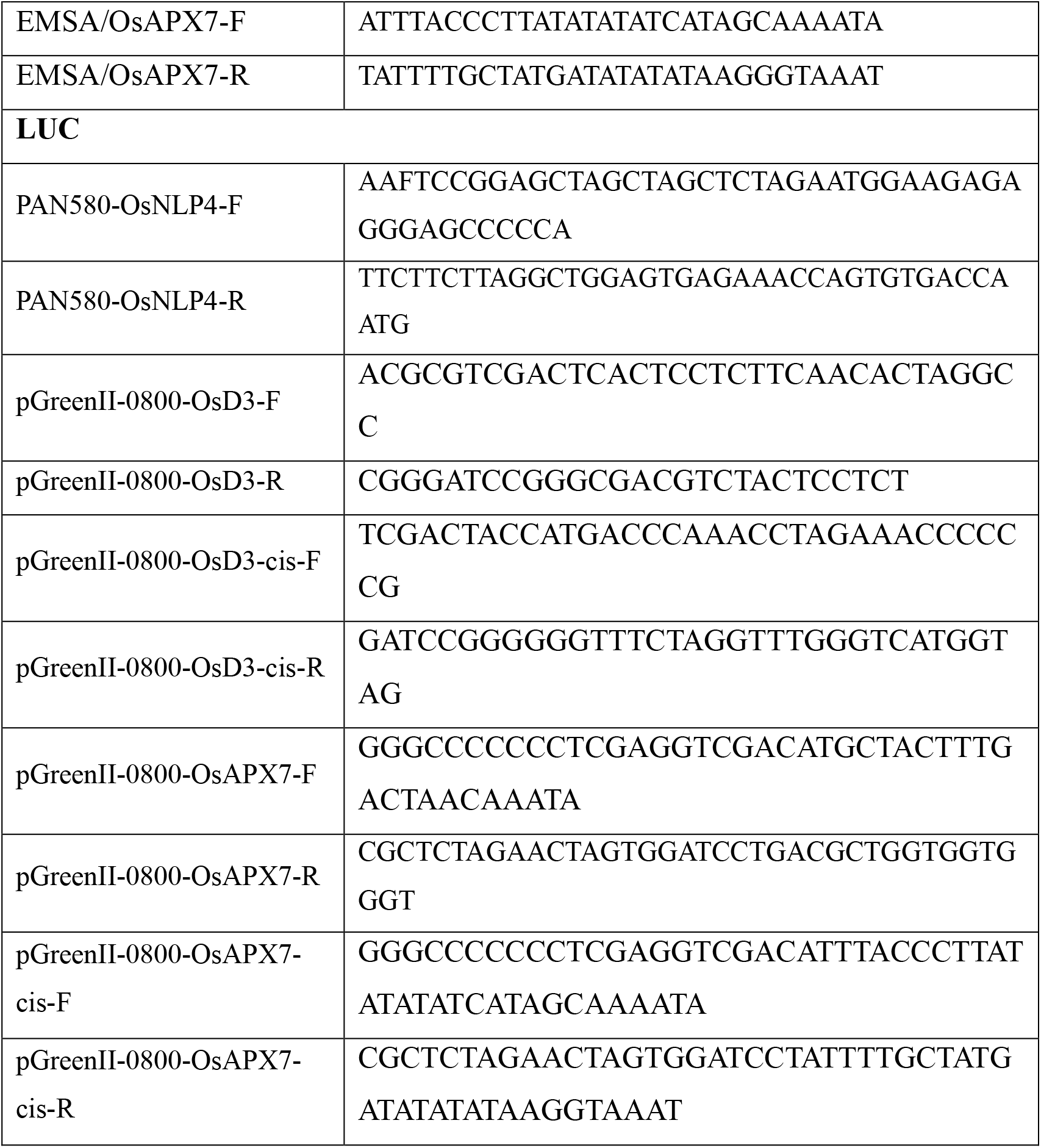
Primers used in this paper.

